# Transcriptome sequencing and screening of genes related to sex determination of *Trichosanthes kirilowii* Maxim

**DOI:** 10.1101/2019.12.17.879635

**Authors:** Xiuqin Hu, Jinkai Zhu, Guanjie Ji, Zhen Wang, Jie Xin

## Abstract

*Trichosanthes kirilowii* Maxim. (*TK*) is a dioecious plant in the Cucurbitaceae for which different sexes have separate medicinal uses. In order to study the genes related to sex determination, transcriptome sequencing was performed on flower buds and leaves of male and female plants using the high-throughput sequencing technology. A total of 145,975 unigenes and 7110 DEGs were obtained. There were 6776 DEGs annotated to 1234 GO terms and enriched to 18 functional groups, including five biological processes related to sugar metabolism. KEGG pathway analysis indicated genes involved in hormone transduction, hormone synthesis and carbohydrate metabolism. The sex determination genes of *TK* are different from known sex determination mechanisms (*ACS11, ACS7(2), WIP1*). Many DEGs of *TK* are involved in reproductive organ formation, hormone signal transduction and regulatory networks. In combination with the previous study of sex differentiation of Cucurbitaceae, the results of GO, KEGG and the expression of related genes in male and female plants, 18 candidate genes for sex determining of *TK* were screened from 151 hormone-related differentially expressed genes. The genes included *MYB80, MYB108* and *MYB21* of the MYB family, *CER1, CBL, ABCB199, SERK1* and *HSP81-3*. The results provide a foundation for the study of sex differentiation in *TK*.

## 1. Introduction

*Trichosanthes kirilowii* Maxim. (namely *TK*) is a perennial climbing herb in the family Cucurbitaceae. Its fruit (fructus trichosanthis), seeds (semen trichosanthis), peel (trichosanthis pericarpium) and root (radix trichosanthis) are all commonly used as traditional Chinese medicines. Due to the large demand for medicinal products in the marketplace, there are many planting bases for *TK* in China. Among these, Changqing District of Jinan City, Shandong Province and the surrounding areas of Feicheng City, Shandong Province have a long history of producing excellent varieties and high-quality medicinal materials that are famous as genuine herbal medicines.

Dioecious plants play an important role in elucidating the mechanism of plant sex determination and evolution, especially plants in the Cucurbitaceae. The studies of sex identification are of great significance in both theory and practice. TK is dioecious and cross pollinated, and the sexes have different medicinal uses. When harvesting seeds and fruits, a large number of female plants (with a small number of male plants) are required, and when harvesting roots, male plants are required. At present, *TK* can be propagated in two ways: vegetative propagation using rhizomes and sexual propagation using seeds. Although the plant sex can be controlled by rhizome propagation, the propagation coefficient is low, and large amounts of raw materials are consumed. Therefore, seed propagation is an economical and practical method of improving the planting efficiency and realizing large-scale cultivation. However, the problem with seed reproduction is that the proportions of male and female plants cannot be controlled. In the natural state, the ratio of males to females is about 7:3. Therefore, it is of great significance to identify the early sexes of *TK* seedlings and to reveal the molecular mechanisms of sex determination.

At present, the methods for sex identification of *TK* include plant appearance, chemical reagents, isoenzymes, protein electrophoresis and molecular markers. Plant sex difference arises from differences in gene expression. Isozymes and proteins are the products of gene expression, and specific gene expression produces specific isozymes or proteins. Therefore, Yu et al. (2003) and Li (2005) used PAGE to identify the early sex of *TK* and found certain differences between male and female strains in enzyme amount and spectral bands. Karmakar et al. (2013) used total proteins of *TK* roots to identify plant sex and found a slight sex difference band with a molecular weight of 19 KDa. Qu et al.’s. (2010) study of RAPD-SCAR have found that S 1200 primers can generate a 600-bp-specific amplification band in the DNA of male *TK*. Guo (2009) carried out isozyme electrophoretic analysis of leaves of *TK* and found that the isozyme bands and enzyme contents from leaves of different sexes were different. While it is generally assumed that sex expression is dominated by the formation and accumulation of flowering substances, the above studies have shown that there is a sex difference in *TK* at the seedling stage.

Extended to the Cucurbitaceae family, although many plants in the family are dioecious, only Concinia indica is identified as having sex chromosomes, that is, an XX/XY sex determination system (Sousa et al. 2013). The sex chromosome of cucumber (*Cucumis sativus* Linn.) is unformed, and the plant sex is controlled by a single gene. Genes related to external hormones, environment, ethylene synthesis and induction of ethylene production are also involved in sex regulation, such as genes CsACS1G and CsACS2 (Boualem et al. 2009; Li et al. 2009). Kumar et al. (2012) found that opc 05-1000 and opc 04-400 can stably amplify different bands in male and female plants of the pointed gourd that can be used for sex identification. Among these, opc 05-1000 bands can mark male plants (i.e., this band only exists in male plants), and opc 04-400 only exists in female plants. The above two markers can be used for sex identification of pointed gourds before flowering. Seneviratme (2002) found that opc-7 can distinguish male and female plants of the pointed gourd; Nanda et al. (2013) found a male linkage marker of the pointed gourd. The research on sex differentiation of *TK* in China is still at the stage of basic exploration, and there are no in-depth reports. In fact, because *TK* is a traditional medicinal plant in China, scientists in other countries know little about it, not only regarding the genome and transcriptome sequencing but also the EST sequence. Although Chinese researchers have done extensive exploratory work in this field and have obtained some basic results, the results, to date, have been insufficient to explore the mechanism of sex determination or find sex determination genes only via the existing sex linkage markers. Therefore, the lack of genomic sequences has become a bottleneck in the study of sex differentiation of *TK*.

The sex differentiation of *TK* lies in the flower organs. The study of flower organ formation will help to understand the morphogenesis and control mechanisms of sex differentiation and will provide a practical basis for the study of the sex genetic law of *TK*.

In this study, we used the leaves and flowers of female and male plants of *TK* to obtain transcriptome information using Illumina sequencing technology. The goals were to search for differential expression genes (DEGs), and to screen for the key genes related to sex differentiation in order to lay a foundation for revealing the sex differentiation mechanism of *TK* at the molecular level.

## 2. Methods and Materials

### 2.1 Sample collection

*TK* is cultivated in the Hebao planting base of Pingyin, Shandong province (116.45 E, 36.28 N). The samples were collected during July 2016. Male and female plants with equal amounts of young leaves, mature leaves, and old leaves were mixed into single samples. The flower buds were collected before, during and after sex differentiation and were mixed into single samples. There were 12 samples with three biological replicates. After sampling, plant leaves and buds were wrapped with tin foil and placed into liquid nitrogen, then stored in an ultra-low-temperature refrigerator.

### 2.2 RNA isolation and quality assessment

Total RNA of each sample was extracted by Tripure, and the concentration and quality were measured using an Aglient 2100 Bioanalyzer, with which RNA integrity (RIN) above 7.5 can be reverse transcribed into cDNA and used to construct transcriptome databases. The database building kit uses NEBNext^®^ Ultra^™^ RNA Library Prep Kit for Illumina^®^.

### 2.3 cDNA library construction, quality control and Illumina sequencing

The constructed library was sent to Beijing Nuohezhiyuan Science and Technology Co., LTD. Using the Illumina HiSeq 2500 sequencing platform for two-terminal sequencing, the size of the sequencing fragment was 151 bp, and the raw sequencing data were obtained.

### 2.4 Raw sequencing processing and de novo assembly

FastQc was used to detect the raw RNA reads and remove the joint sequences. The leaves and flowers were taken as a whole sample. Trinity was used for de novo assembly, and over loop was used to splice the Contig and Unigene fragments of clean reads to obtain the Unigenes.

### 2.5 Screening of differentially expressed genes

Cuffdiff of Cufflinks software was used to analyze the differences in gene expression levels in each group to identify the differentially expressed genes (DEGs). Cuffdiff uses non-parametric statistical methods to estimate the mean and variance of FPKM values in different samples based on annotation files and identifies selected transcripts with significant differences in expression between samples through *t* tests.

### 2.6 Functional annotation and classification

Through BLAST comparison software, the unigenes sequences were compared with the protein databases Nr and KEGG. Classification information and gene function annotation were carried out by BLASTx. In order to reflect the expression of sex difference genes more accurately, the GO function and KEGG pathway significance enrichment analyses were carried out to determine the main biological functions and the main metabolic pathways that the genes were involved in. GO enrichment analysis was performed on DEGs using the SEA tool of agriGo software, and the *P* values were statistically analyzed and corrected (FDR ≤ 0.05) using Fisher’s exact test and the Bonferroni correction method. The KEGG pathway enrichment analysis uses KOBAS (KEGG Orthology-based Annotation System, http://kobas.cbi.pku.edu.cn/home.do), where the calculation principle is the same as in the GO function enrichment analysis. To control the false positive rate, BH (Benjamini and Hochberg’s test) was used for multiple tests with *P* = 0.05. A KEGG pathway meeting the above conditions was defined as a significantly enriched pathway.

## 3 Results and analysis

### 3.1 Transcriptome sequencing and de novo assembly

We constructed 4 transcriptome libraries: female flower buds, female plant leaves, male flower buds, and male plant leaves. The four libraries respectively obtained 16545286, 17914519, 16699544 and 17619567 high-quality reads. The effective detection rate of each library was above 90%. Transcriptome splicing was performed taking leaves and flowers as a whole unit. For species without reference genomes, de novo assembly is the most commonly used technique, and thus Trinity software was used to assemble the sequencing data. In all, 145,975 Unigene fragments were obtained after redundancy was removed. Specific assembly results are shown in Table 1. Among the resulting fragments, 400–800 nt had 13,305 pieces; 400–800 nt had 3,643 pieces; 1600–2000 nt had 1931 pieces; 2200 and 2600 nt had 919 pieces; 2800–3200 nt had 485 pieces; 3400–3800 nt had 248 pieces and there were 879 pieces ≥ 4000 nt.

**Table 1.**
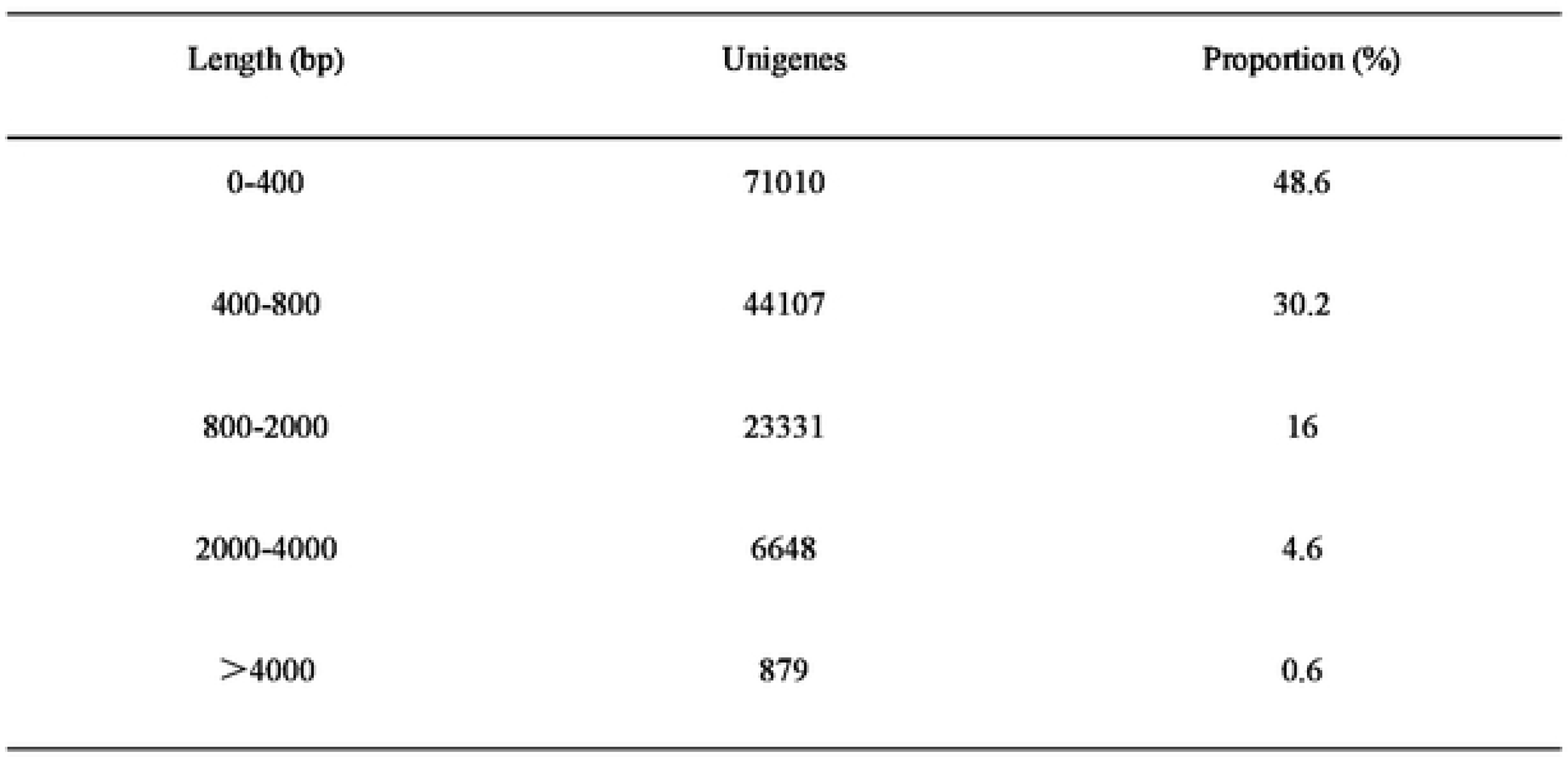
Length distribution of assembled Unigenes from *TK*

| Length (bp) | Unigenes | Proportion (%) |
| --- | --- | --- |
| 0-400 | 71010 | 48.6 |
| 400-800 | 44107 | 30.2 |
| 800-2000 | 23331 | 16 |
| 2000-4000 | 6648 | 4.6 |
| >4000 | 879 | 0.6 |

### 3.2 Identification and analysis of DEGs

Cufflinks was used to quantify the expression of DEGs, and the FPKM value reflected the expression level of each gene. FPKM ≥ 10 was set as a highly expressed gene, 2 ≤ FPKM < 10 as a medium expression gene, and FPKM < 2 as a low expression gene. According to the FPKM standard, we calculated the gene expression levels of flower bud samples as follows: the number of low expression genes in female flower buds was 2491, and the number of medium expression genes was 1924; the number of high expression genes was 1165; the number of low expression genes in male flower buds was 1588; the number of medium expression genes was 2463, and the number of high expression genes was 1530. There were fewer low expression genes in male flower buds than in female flower buds, but more genes were expressed in male flower buds than in female plants. There were few differences in the medium expression genes of male and female flower buds, but there was a large difference in the number of high expression genes. Then, we used Cuffdiff to calculate the significance of differential gene expression, and set fold change > 2 or < 0.5, *P* < 0.01 as the marker for identifying DEGs. We defined the up-regulated genes in female flower buds as up-regulated genes; in contrast, down-regulated genes in male flower buds were defined as down-regulated genes. The statistics of DEGs are shown in Table 2. The number of DEGs in leaves of male and female plants was 1056; the number of up-regulated genes was 590, and the number of down-regulated genes was 466. The number of DEGs in flower buds was 5580; the number of up-regulated genes was 3104, and the number of down-regulated genes was 2476.

**Table 2.** Statistical analysis of differentially expressed genes in flower buds and leaves of *TK*

| Analysis | DEGs | UP | DOWN |
| --- | --- | --- | --- |
| flowers | 5580 | 3104 | 2476 |
| leaf | 1056 | 590 | 466 |

### 3.3 Functional annotation and classification of DEGs

All the differentially expressed sequences were submitted to NCBI for BLASTn comparison. A total of 5303 unigenes had annotations in the NR databases. Of those unigenes, 74% (3924) obtained homologous genes or obtained gene notes, and 26% (1379) had no homologous sequences or were position genes without functional annotation, as shown in Fig. 1. In the near source species that Unigenes matched in the NR database, cucumber accounted for the highest proportion (15184, 44.95%), followed by *Cucumis sativus* (12117, 35.87%), *Vitis vinifera* (779, 2.31%), Arabidopsis (455, 1.35%), *Citrus sinensis* (387, 1.15%), *Cucumis melo* subsp. Melo (384, 1.14%), and other species (15.52%); 82% of the genes were annotated to Cucurbitaceae (Fig. 2).

**Fig. 1.**
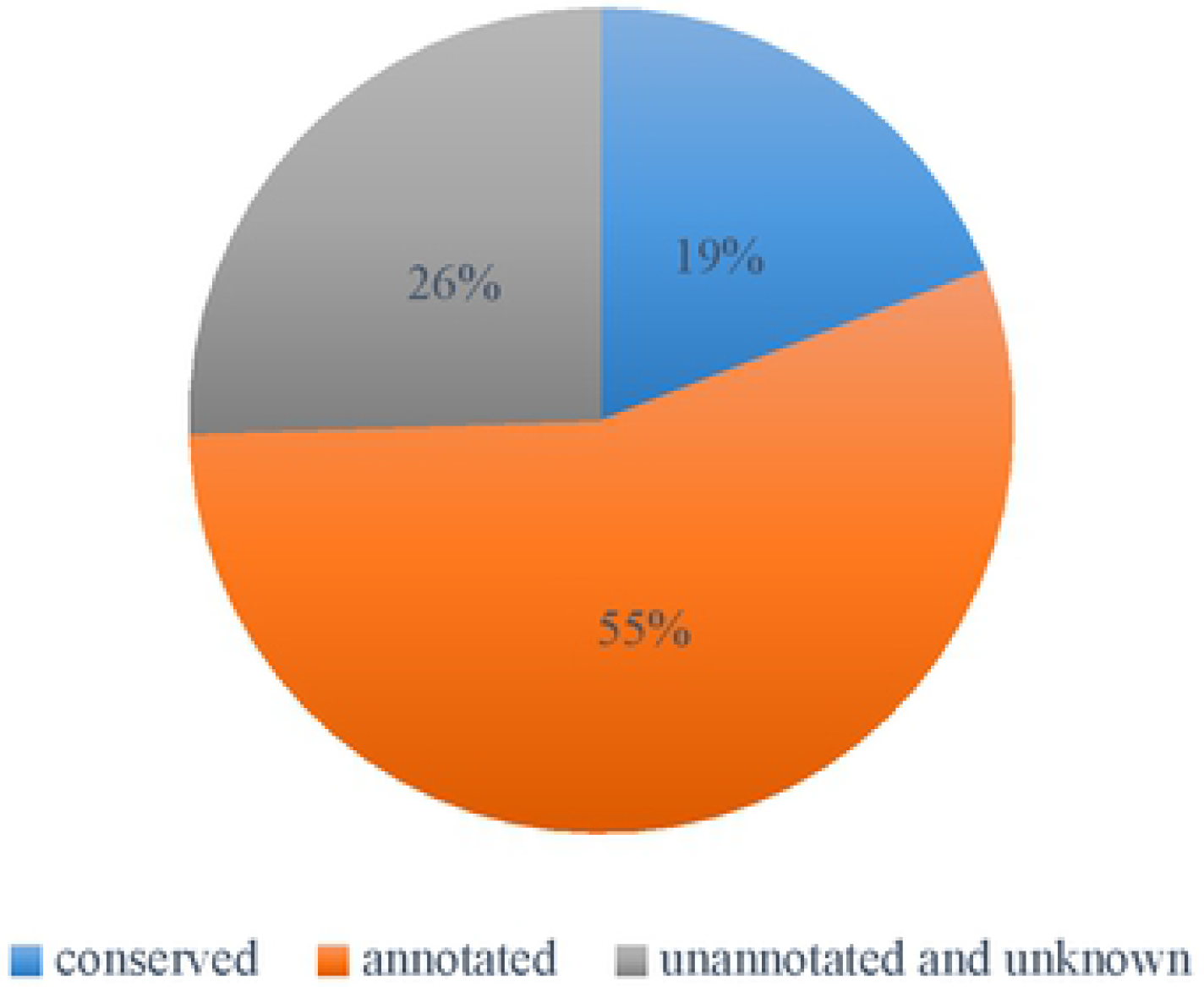
Functional annotation of the differential expression Unigenes of *TK* in the nonredundant databases

**Fig. 2.**
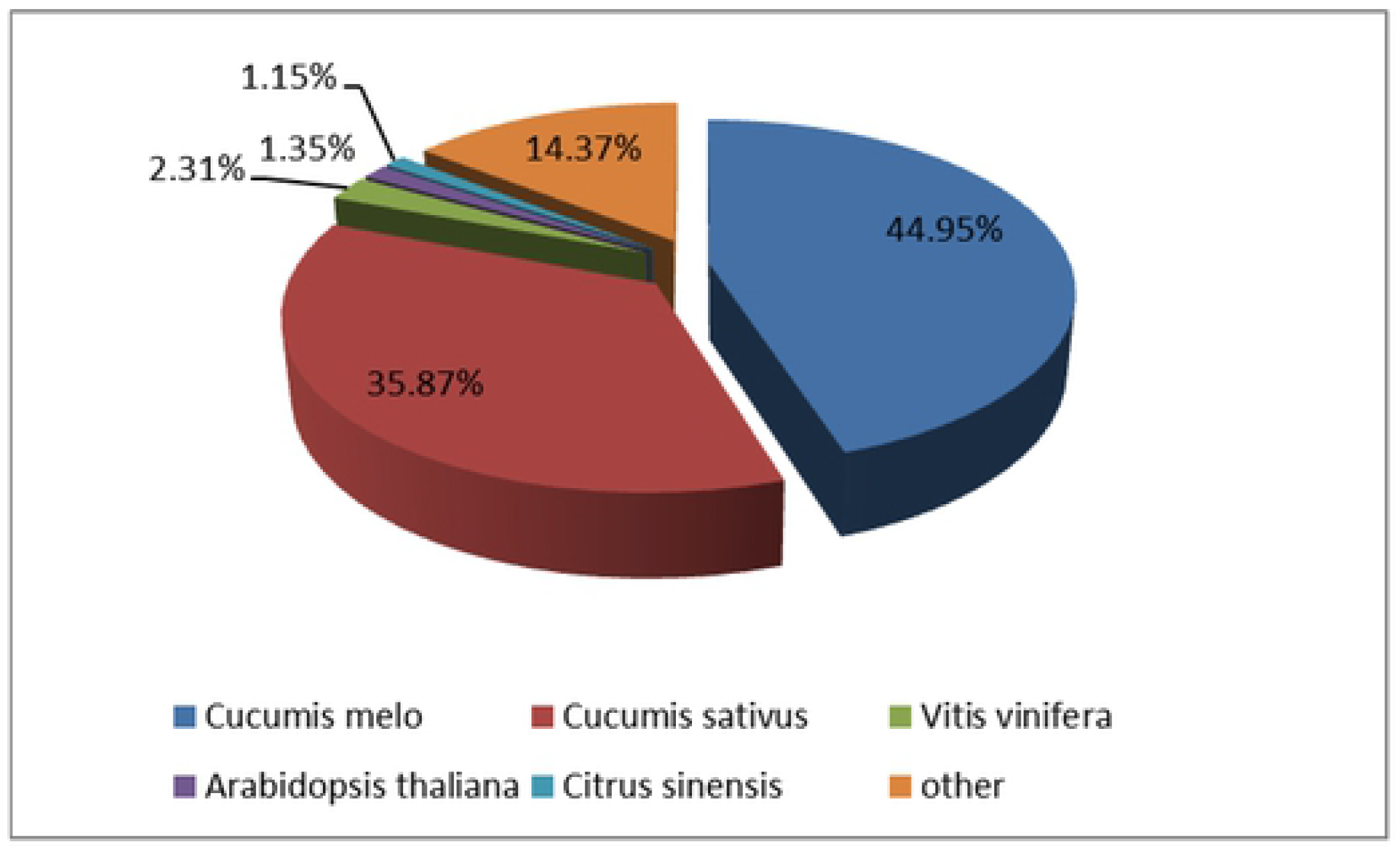
Annotation and classification of the assembled unigenes in the nonredundant databases, and the species distribution of BLAST hits for each unigene in the Nr database

### 3.4 Analysis and functional classification of DEGs

In order to further clarify the function of differentially expressed genes, the selected DEGs were analyzed with Blast2GO. The software comprehensively considers the similarity of target sequences and alignment sequences, GO item source reliability, and the structure of a GO directed acyclic graph, and extracts the qualified GO functional items in the mapping process (GO terms) annotated to the target protein (DEG protein). In this study, 6776 DEGs annotated 1234 GO items, including cell components (133), molecular functions (353) and biological processes (748). All the matched gene sequences were further enriched into 46 functional categories, among which the functional groups of membrane part, membrane, binding, catalytic activity, cellular process and metallic process contained more unigenes, while biological adhesions, location, protein binding, growth, extractor region part, rural reservoir activity and immune system contained fewer unigenes (Fig. 3). Then according to the GO annotation information of significantly differentially expressed genes, we further analyzed the significance of enrichment and calculated *P* values by Fisher’s exact test (FET). If FDR ≤ 0.05 and FDR ≤ 0.01, we assumed that there was significant enrichment or extremely significant enrichment of this GO function. The differentially expressed genes were enriched in 18 functional groups. These genes included the cell wall polysaccharide metabolic process (GO:0010383), hemicellulose metabolic process (GO:0010410), xyloglucan metabolic process (GO:0010411), hydrolase activity, acting on glycosyl bonds (GO:0016798), and xyloglucan: xyloglucosyl transferase activity (GO:0016762). The complete results are listed in Table 3.

**Fig. 3.**
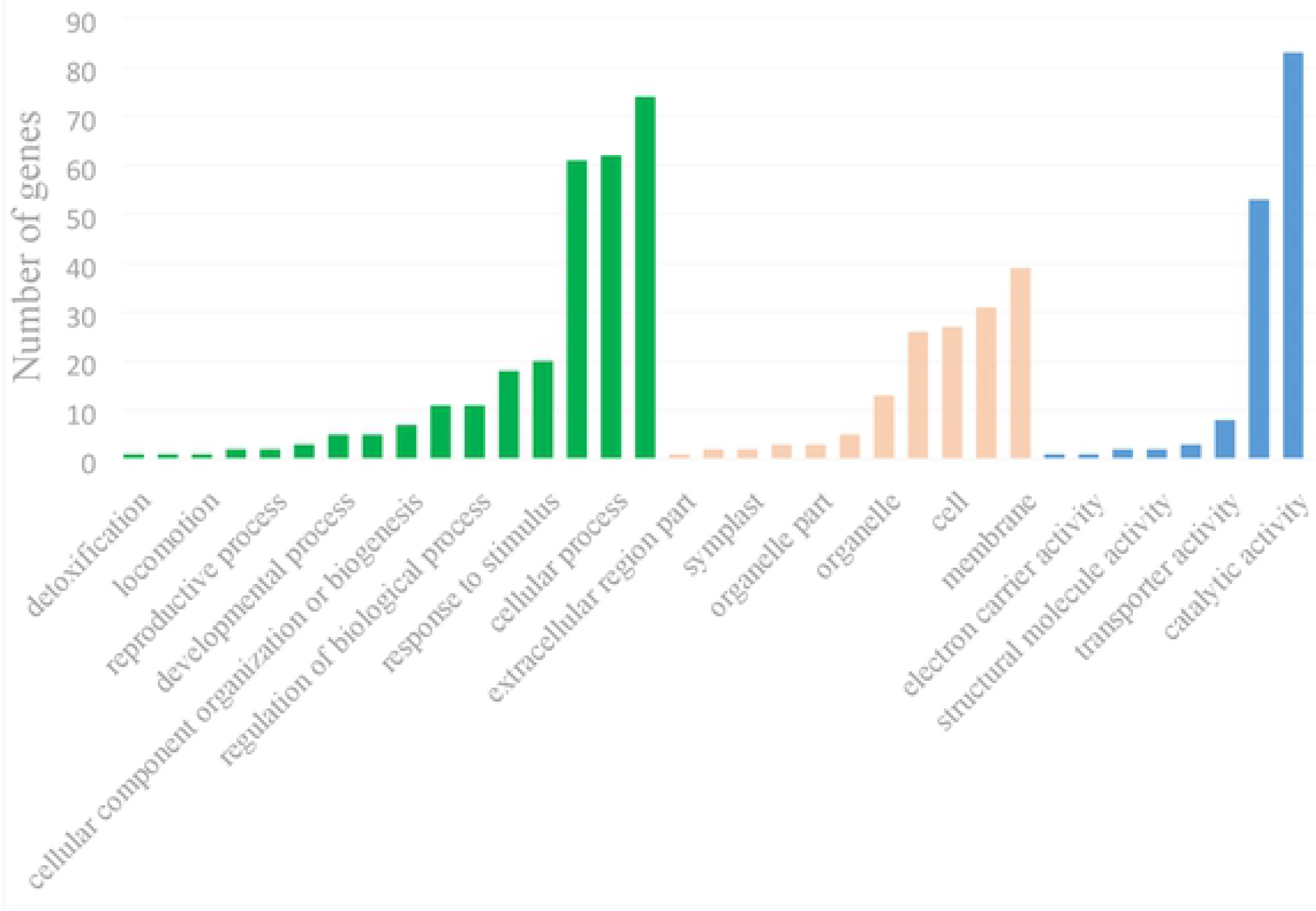
Gene ontology categories of the assembled unigenes from the *TK* Unigenes were assigned to three categories: cellular components, molecular functions and biological processes

**Table 3.**
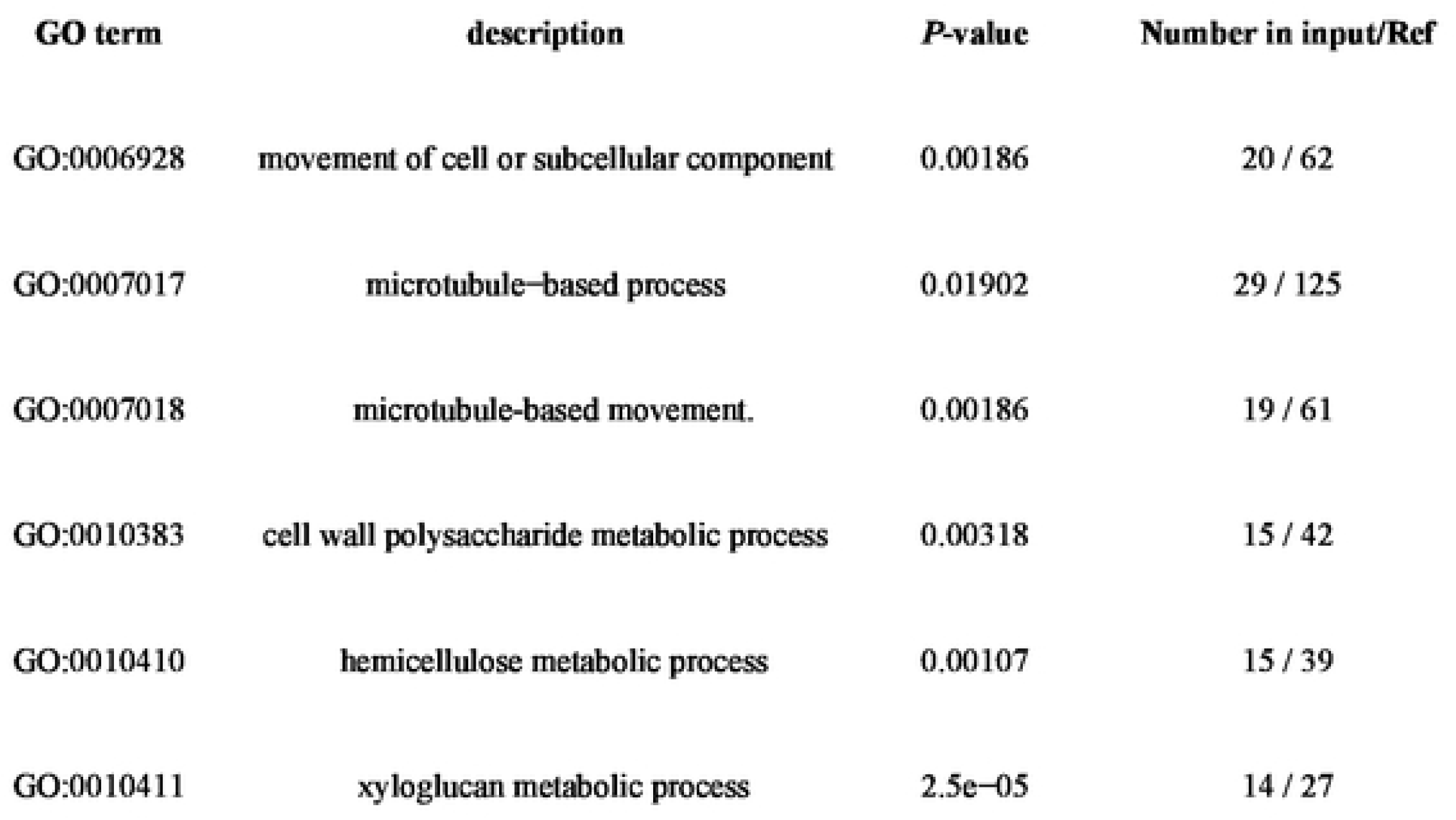
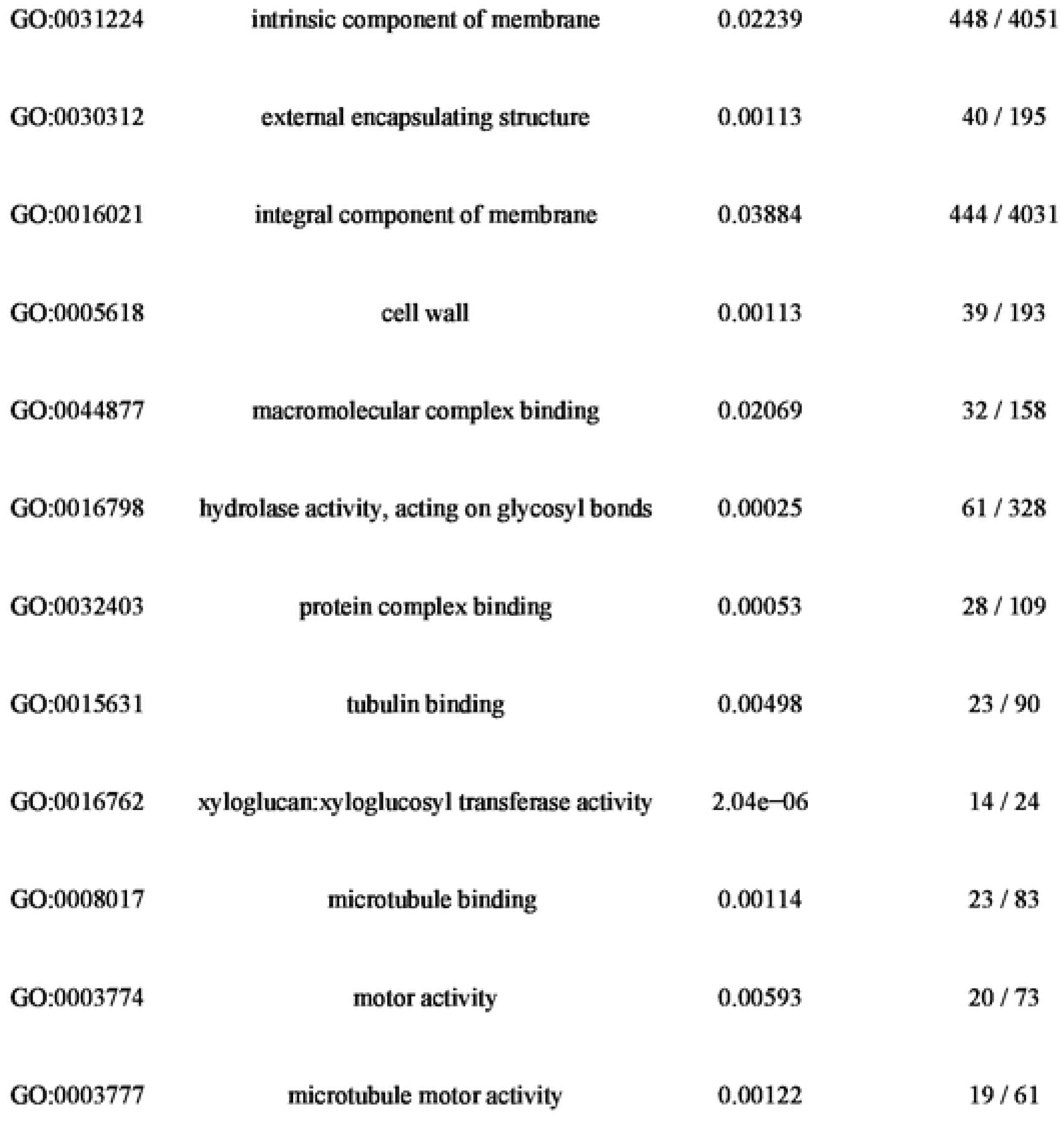
Significant enrichment of differentially expressed genes by GO

In order to further elaborate the biochemical pathways expressed by differential expression genes, KOBAS was used to compare the differential expression genes to the plant KEGG database, and an *E* value < 10^-5^ was set to identify the possible biological pathways. A total of 2286 different genes were located in 131 pathways, as shown in Fig. 4. The pathways with more genes included global and overview maps, translation, carbohydrate metadata, environmental adaptation, folding, sorting and graduation, while endocrine and metallic diseases only had one gene. Twenty pathways were significant (*P* < 0.05; Table 4).

**Fig. 4.**
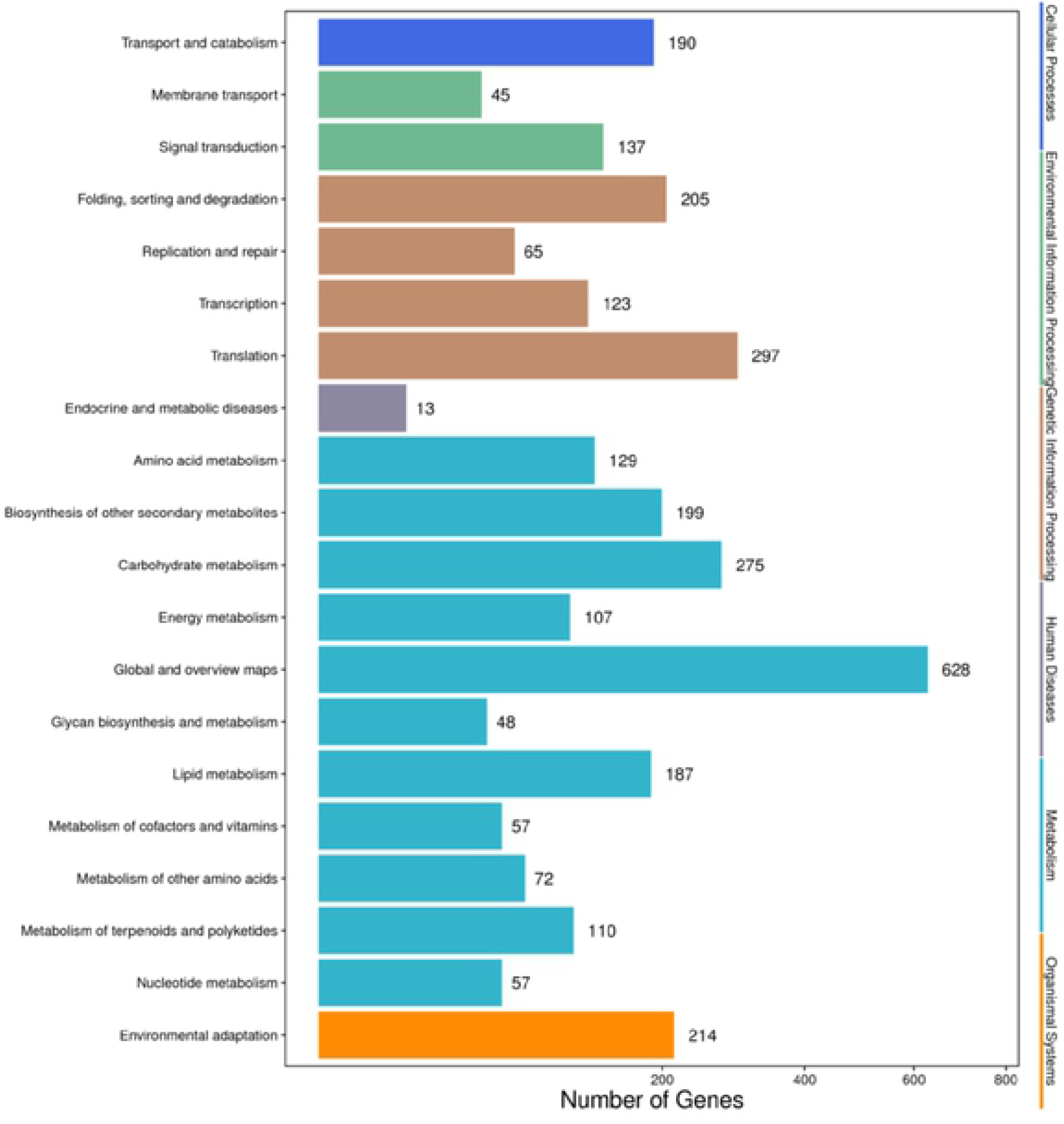
Distribution of Kyoto Encyclopedia of Gene and Genomes (KEGG) pathways in *TK*

**Table 4.** Significant difference enrichment pathway by KEGG

| No. | Pathway | KO ID | P-Value |
| --- | --- | --- | --- |
| 1 | Linoleic acid metabolism | Ko 00591 | 0.00000 |
| 2 | Phenylpropanoid biosynthesis | Ko 00940 | 0.00000 |
| 3 | Stilbenoid, diarylheptanoid and gingerol biosynthesis | Ko 00945 | 0.00000 |
| 4 | Flavonoid biosynthesis | Ko 00941 | 0.00000 |
| 5 | Biosynthesis of secondary metabolites | Ko 01110 | 0.00003 |
| 6 | Isoflavonoid biosynthesis | Ko 00943 | 0.00010 |
| 7 | Plant hormone signal transduction | Ko 04075 | 0.00013 |
| 8 | Limonene and pinene degradation | Ko 00903 | 0.00047 |
| 9 | Cyanoamino acid metabolism | Ko 00460 | 0.00132 |
| 10 | Plant-pathogen interaction | Ko 04626 | 0.00245 |
| 11 | Pyruvate metabolism | Ko 00620 | 0.00460 |
| 12 | Ribosome | Ko 03010 | 0.00485 |
| 13 | Ascorbate and aldarate metabolism | Ko 00053 | 0.00788 |
| 14 | Steroid biosynthesis | Ko 00100 | 0.01007 |
| 15 | mRNA surveillance pathway | Ko 03015 | 0.01347 |
| 16 | Circadian rhythm - plant | Ko 04712 | 0.01441 |
| 17 | RNA polymerase | Ko 03020 | 0.02521 |
| 18 | Glycolysis / Gluconeogenesis | Ko 00010 | 0.02724 |
| 19 | Caffeine metabolism | Ko 00232 | 0.04228 |
| 20 | Pentose and glucuronate interconversions | Ko 00040 | 0.04994 |

### 3.4 Screening of genes related to sex differentiation

The sex of cucumbers and melons in the Cucurbitaceae is controlled by the *ACS11, ACS7(2*) and *WIP1* genes (Saito S et al. 2007; Wu T et al. 2010; Boualem A et al. 2015). The homology of *CmACS2* and *CsACS7* reached 96.19%, and the homology of *CmACS11* and *CsACS11* reached 92.52%, indicating that the two genes are highly conserved, but these three sex determining genes were not found by the primer amplification and transcriptome data match in *TK* (the primer design is shown in Table 5). Referring to the mechanism of sex differentiation in other Cucurbitaceae plants, we continued to believe that hormone genes or genes induced by hormones are the main factors determining sex differentiation of *TK*. Genes related to ethylene synthesis and genes induced by ethylene are particularly important (Shannon S and De La Guardia MD 1969; Rudich J et al. 1972; Takahashi H and Jaffe MJ 1984; Trebitsh T et al. 1987; Yamasaki S et al. 2001; Yamasaki S et al. 2003). We analyzed the DEGs related to hormones in *TK* by using blast P. A total of 7110 differential genes were compared with the Arabidopsis hormone database, and when the *E* value < 10^-6^ or the similarity ≥ 60%, we considered the two proteins to be homologous. According to this standard, we found 151 genes related to hormones from the DEGs, including 19 genes related to hormone synthesis, three genes related to hormone metabolism, 6 genes related to hormone receptors, 14 genes related to hormone response, 91 genes related to hormone signal transportation, and 18 genes related to hormone transportation (Fig. 5). In combination with the studies of sex differentiation of Cucurbitaceae, the results of GO, KEGG and gene expression patterns in male and female plants, 18 candidate genes for sex differentiation of *TK* were screened; these are listed in Table 6.

**Fig. 5.**
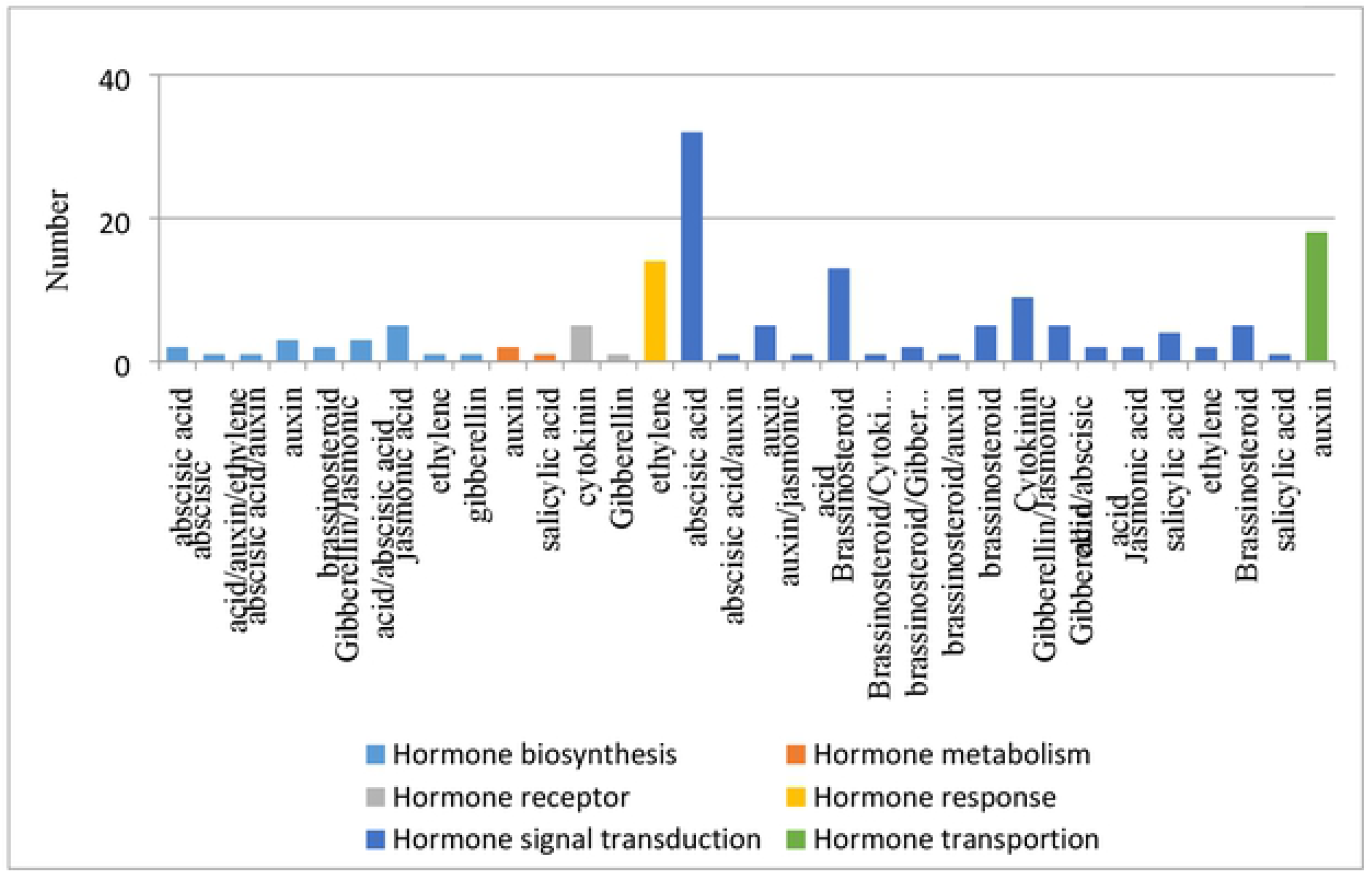
Hormone-related genes in *TK*

**Table 5.**
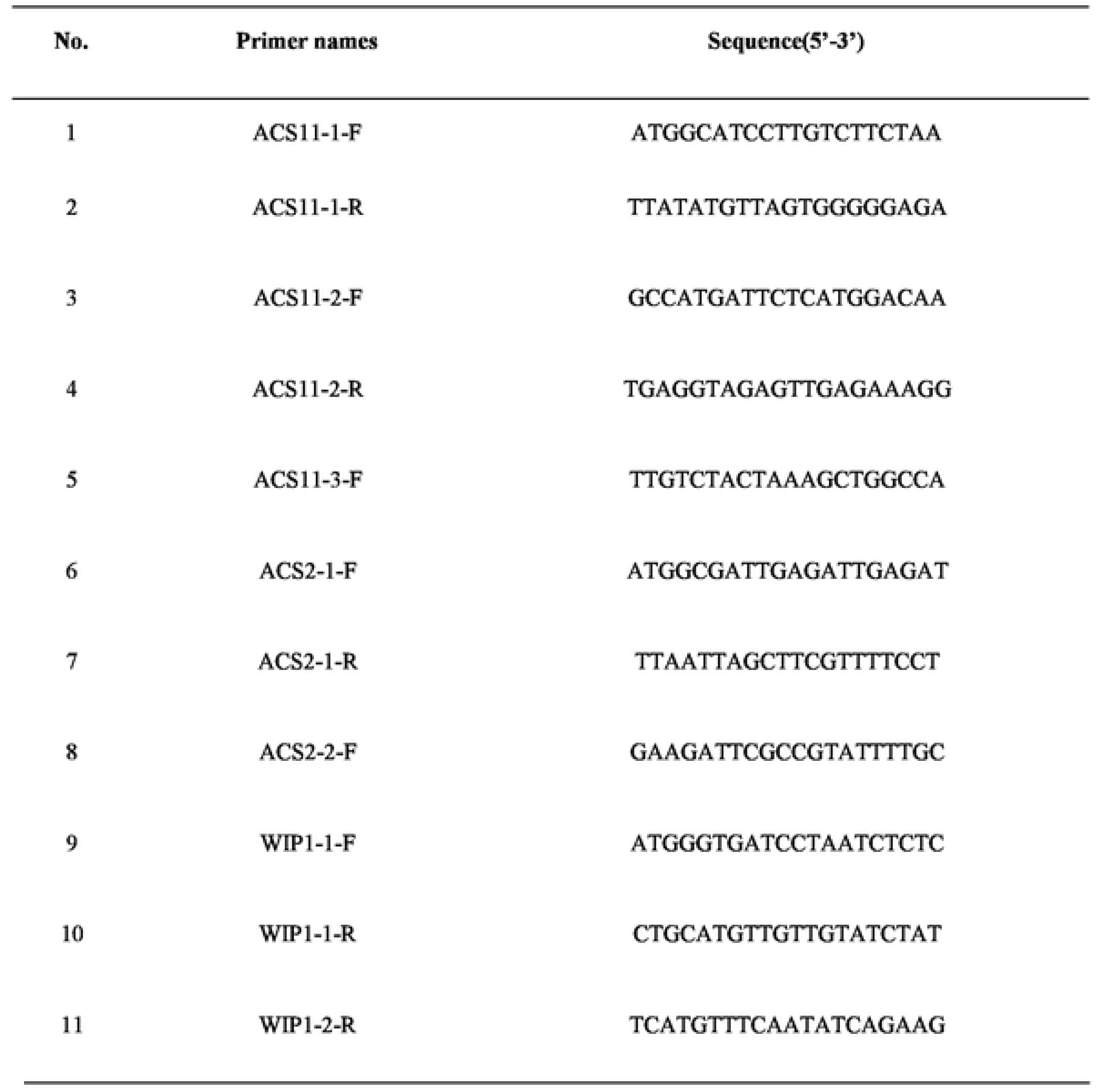
The design of primers for amplification of candidate genes

**Table 6.**
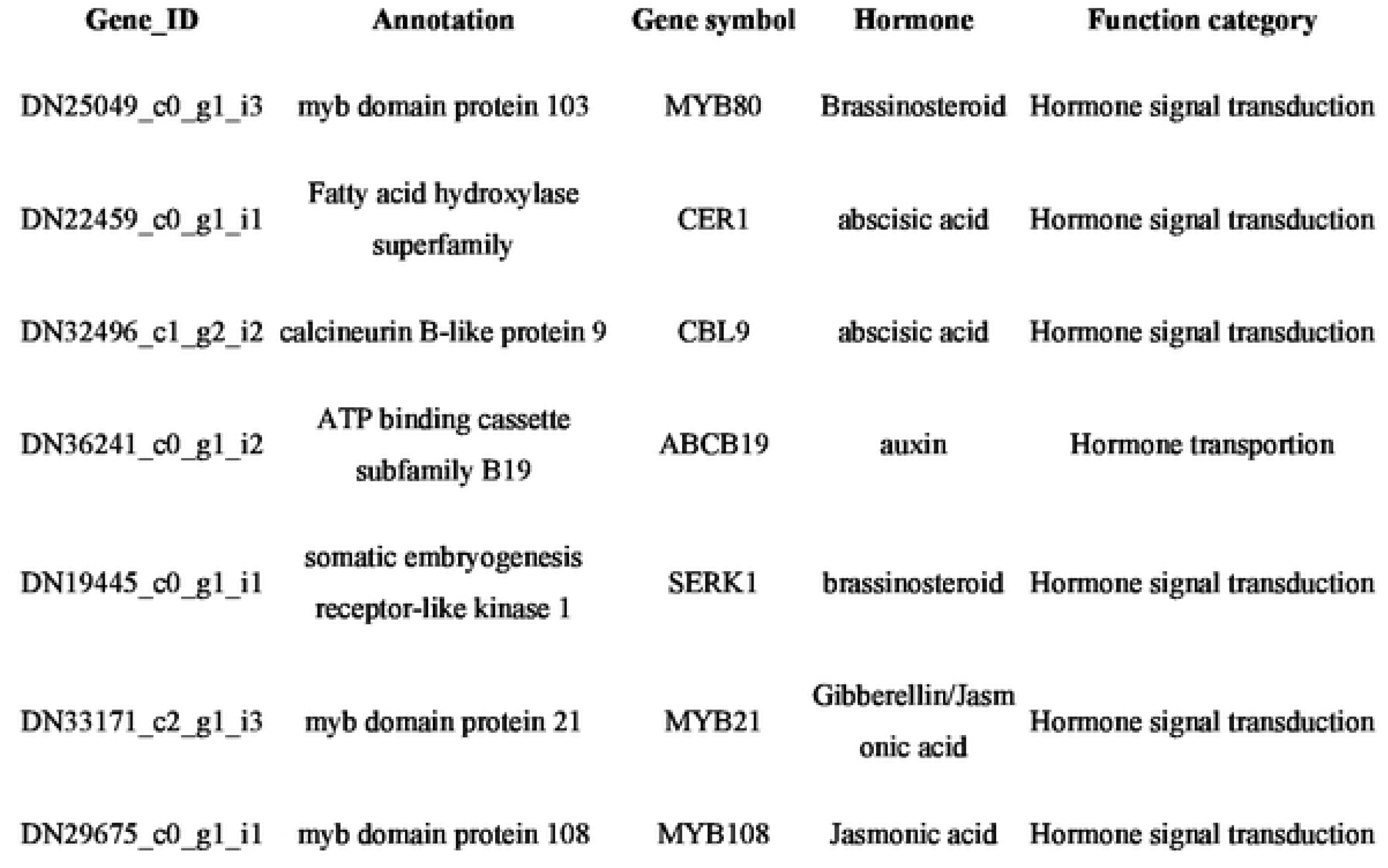
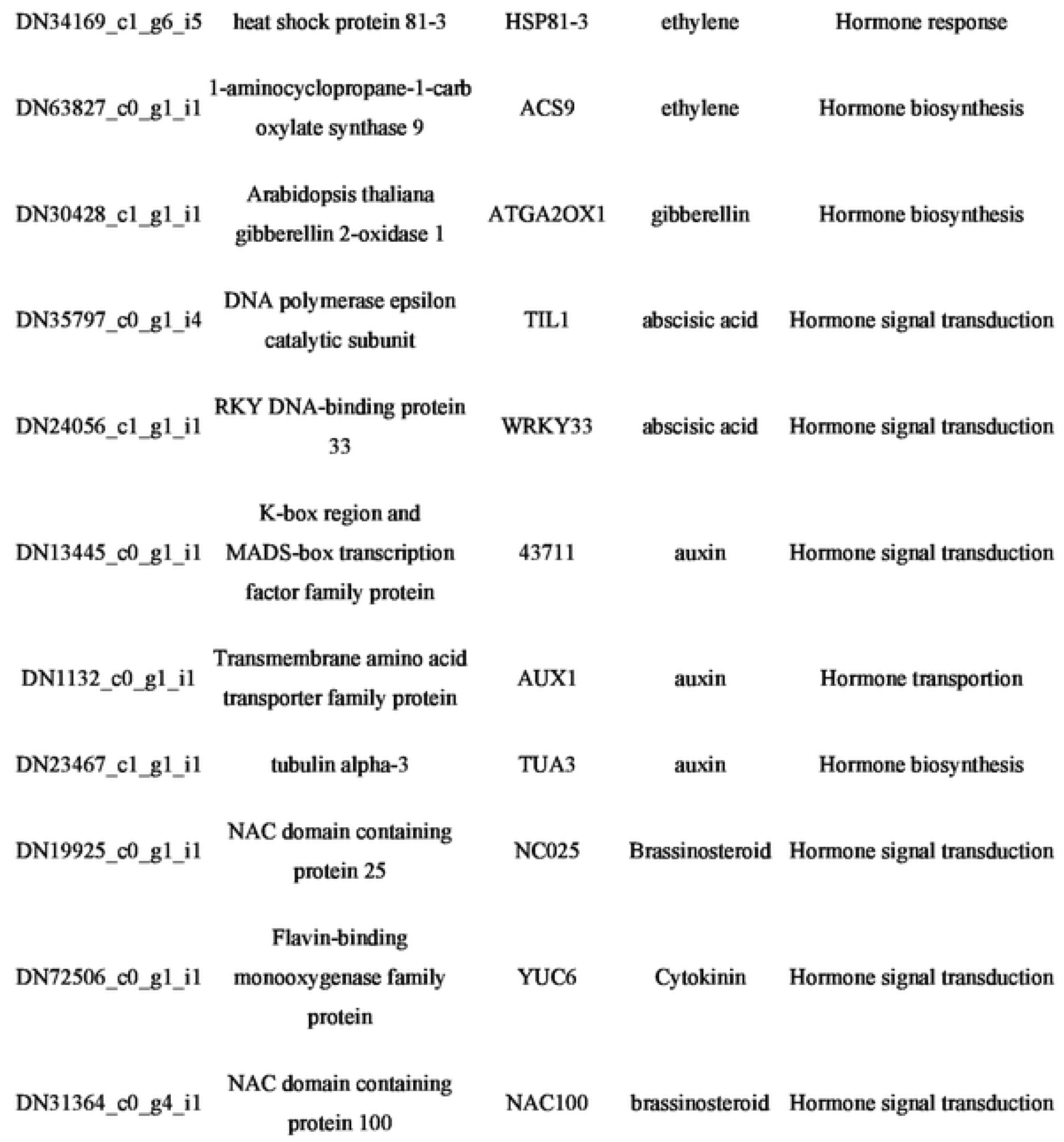
Candidate genes related to sex differentiation of *TK*

## 4. Conclusion

Plant sex determination and differentiation have become a major focus of developmental genetic research in recent years. Compared with animals, plants have more variable sex determination patterns. Stamens and carpels require a large number of specific genes to participate in each development stage. Cucurbitaceae species are numerous, and their sexual systems are also variable. For example, the flower primordium of cucumbers is bisexual at first, and then the stamen or carpel stops development selectively, forming a unisexual flower (Bai S L et al. 2004). However, the female flowers of *TK* are bisexual initially and the stamen development then stops, but the male flowers of *TK* are completely unisexual (Xin J et al. 2018). In addition, hormones and environmental factors can affect sexual development in the Cucurbitaceae, and ETH plays a major role (Wu T et al. 2010). For example, using ETH on monoecious watermelon plants will change all flowers into female flowers. In contrast, treatment of watermelon female plants with ETH inhibitors will lead to the occurrence of bisexual flowers. Consistent with the fact that ETH is a female hormone, watermelon gene A and cucumber gene M, as homologous genes, both encode the rate-limiting enzyme ACS during the ETH synthesis.

Our aim was to discover the sex determining genes in *TK*. The male and female flower buds and leaves of *TK* before, during and after sex differentiation were selected as research materials according to the previous study (Xin J et al. 2018). After screening, 7110 differentially expressed genes were obtained, including 3694 up-regulated genes and 2942 down-regulated genes. Many genes involved in the formation of reproductive organs, hormone signal transduction and regulatory networks were indicated. In all, 6776 DEGs were annotated to 1234 GO items, and GO was enriched in 18 functional groups, including five biological processes related to carbohydrate metabolism: cell wall polysaccharide metabolic process (GO:0010383), hemicellulose metabolic process (GO:0010410), xyloglucan metabolic process (GO:0010411), hydrolase activity, acting on glycosyl bonds (GO:0016798), and xyloglucan: xyloglucosyl transferase activity (GO:0016762). This indicates that carbohydrate metabolism plays an important role in the sex differentiation of flower buds. Based on the KEGG pathway analysis, different genes of male and female plants were significantly enriched in regard to phylopanoid biosynthesis (KO 00940), stilbenid, dialheptanoid and ginger biosynthesis (KO 00945), biosynthesis of secondary metals (KO 01110), plant hormone signal transmission (KO 04075), limonene and pinene degradation (KO 00903), cyanoamino acid metabolism (KO 00460), steroid biosynthesis (KO 00100), RNA polymerase (KO 03020), glycolysis / glycogenesis (KO 00010), pentose and glycornate conversions (KO 00040); this suggest that hormones and sugars may be involved in the sex differentiation of *TK*. In view of the effect of hormones on the sex of Cucurbitaceae plants, we carefully analyzed the gene expression of HRGs. Combined with the results of GO and KEGG, 18 candidate genes for sex determination were screened from 151 hormone-related differential genes. These included the MYB family of *MYB80, MYB108* and *MYB21, CER1, CBL, ABCB199, SERK1, HSP81-3* (Aarts MG et al. 1995; Mandaokar A and Browse J 2009; Phan HA et al. 2011; Phan HA et al. 2011; Song SS et al. 2011; Mähs A et al. 2013; Xu Y et al. 2014; Cecchetti V et al. 2015). The results will provide a foundation for the study of the sex differentiation mechanism of *TK*.

## Acknowledgments

This work was supported by the foundation of National Natural Science Foundation (Nos. 81803651 and 31701889) and Shandong Provincial Natural Science Foundation (No. ZR2019PH088). We thank LetPub (www.letpub.com) for its linguistic assistance during the preparation of this manuscript.

